# Consistent long-distance foraging flights across years and seasons at colony level in a Neotropical bat

**DOI:** 10.1101/2024.07.29.605243

**Authors:** María C. Calderón-Capote, Mariëlle L. van Toor, M. Teague O’Mara, Travis D. Bayer, Margaret C. Crofoot, Dina K.N. Dechmann

## Abstract

All foraging animals face a trade-off: how much time should they invest in exploitation of known resources versus exploration to discover new resources? For group-living central place foragers, balancing these competing goals poses particular challenges. The availability of social information may discourage individuals from investing in risky, expensive but possibly rewarding exploration. We GPS-tracked groups of greater spear-nosed bats (*Phyllostomus hastatus*) from three colonies on Isla Colón in Panamá. In the dry season, when these omnivores forage on the nectar of ephemeral balsa flowers (*Ochroma pyramidale*), bats consistently travelled long distances to remote, colony-specific foraging areas, bypassing flowering trees closer to their roosts. They continued to use these same areas in the wet season, when feeding on a diverse, presumably ubiquitously distributed diet, but also visited other, similarly distant foraging areas. Foraging areas were shared within, but not always between colonies. Our longitudinal dataset suggests that bats from each colony invest in long-distance commutes to socially learned shared foraging areas, bypassing other available food patches. Rather than investing in exploration to find nearby resources or engaging in a win-stay lost-shift foraging strategy, these bats follow colony specific behaviours consistent with the existence of culturally transmitted preferences for specific feeding grounds.

## Introduction

Foraging is vital and direct determinant of organismal fitness. Foraging animals have to maintain a delicate balance between exploitation and exploration [1–5]. They must weigh the decision to exploit known resources against the potential benefits of seeking out new ones [2,6]. This balance hinges on three main factors: environmental conditions (i.e., quality and quantity of available resources), individual traits (prior information, cognitive abilities) and social interactions. Social central place foragers often forage in the presence of others and can learn from them. For example, social information can deter risky exploration while encouraging exploitation [1,7,8]. Understanding how animals navigate this trade-off is essential for uncovering group dynamics and the development of potential social traditions.

Greater spear-nosed bats (*Phyllostomus hastatus*) are omnivores described to forage within <10km of their roost in Trinidad [9,10]. In the dry season, they forage socially on the nectar of ephemeral balsa trees. GPS tracking in Panamá revealed *P. hastatus* flying individually >25 km to their foraging areas when blooming balsa were particularly scarce [11]. This intraspecific variation offers the opportunity to investigate how social foraging may be mediated by the resource landscape and how this results in a trade-off between exploration and exploitation. We tracked foraging *P. hastatus* over six-years in three colonies during the dry and wet season in Isla Colón, Panamá. Based on the literature we predicted that 1) bats should forage on balsa together and within 10 km during regular dry seasons. We anticipated increased exploration (increased tortuous movements) during the wet season, when bats switch to more evenly distributed insects and fruit. We expected bats to forage alone and closer to the roost in the wet season. 2) We predicted colonies would use separate foraging areas at least during the dry season to avoid competition for flowering balsa trees. 3) Finally, we expected switching foraging areas between seasons, reflecting shifts in resource availability and preferences. We instead found a pattern of long-distance travel to largely consistent foraging areas, and developed a set of movement simulations to test whether the spatial distribution of foraging sites we observed diverged from the expectations. The results of this study will help understand how group living animals adjust their foraging decisions to the resource availability and knowledge the of local landscape.

## Methods

### Tracking *Phyllostomus hastatus* movements

We captured 216 individuals (134 females, 82 males) at three different colonies on Isla Colón, Bocas del Toro, Panamá, during the dry (February-March) in 2016 and 2022, and wet season (December, August) in 2021 and 2023. Colony 1, located at the centre of the island, and Colony 2, located 5 km away at the northernmost tip of the island, are each home to ∼500 bats. Colony 3, located 1.2 km south of Colony 1, has a population of ∼150 bats (Figure 1).

**Figure 1.**
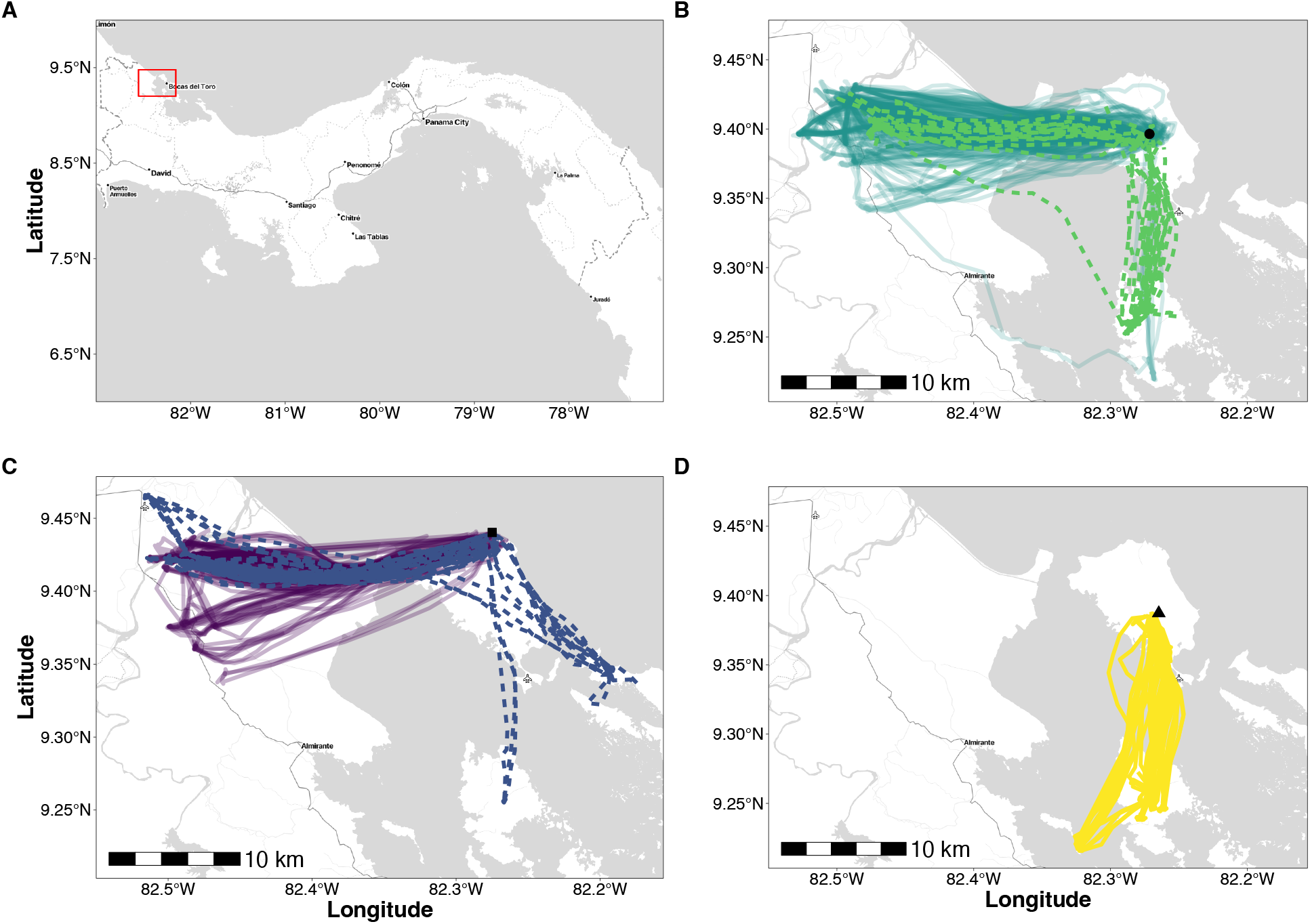
Consistent, colony-specific long-distance foraging flights across years and seasons. A) Map of Panama, inset: study area. B) colony 1 (wet and dry seasons 2016-2023). C) colony 2 in March 2022 (dry) and August 2023 (wet). D) colony 3 in March 2022 (dry). Roosts: circle = colony 1; square = colony 2; triangle = colony 3. Dotted lines: wet season, solid lines: dry season.

We caught bats using a ring trap placed over the roosting cavities. We determined sex, reproductive status and age, measured forearm length (± 0.01 mm), mass (± 0.5 g), and marked them with subcutaneous PIT tags (ID 100 Transponder, Trovan^®^). We tracked only adults with different biologgers and programming schedules (Table S1). Tags were wrapped in shrink tube that we glued (Osto-bond, Montreal Ostomy) to the back of the bats [11]. Biologgers weighed 6.53 ± 0.49 g and represented 5.36 ± 4.82 % of bat mass. Females were in early pregnancy in March 2022, but did not lose substantial weight (pre-tagging mass (n = 20): 109.15 ± 15.46 g; post-tagging = 105.6 ± 12.77 g). Tag recovery varied across colonies and years (Table S2). Tags collected data from 18h–06 h local time.

### Movement analysis

GPS outliers and points with speeds >15 m/s (unlikely for this species) were removed from the data. We down-sampled GPS data to two- (or in March 2022 three-) minute intervals to correct for different sampling rates (Table S1). We used only complete individual tracking nights to calculate distances, bearings and activity (Table S2). From tracks which missed out- or inbound commutes, we only calculated mean distances, directions and in the shared foraging distance/angle analysis (simulations below).

### Behavioural classification

We fitted a three-state hidden Markov model (HMM) for each bat night using the momentuHMM package to identify behaviours [12]. To implement the HMM we first regularised the tracks by inserting “NA” for missing observations to obtain a complete series of two- or three-minute intervals, using the setNA function from the adehabitatLT package [13]. A previous study found social resting between foraging as an important behaviour [11]. However, after down-sampling the data resolution did not accurately allow to distinguish between the categories used there (slow/fast foraging and resting). Thus, we fitted a two-state model with “foraging” (short movements with low persistence of direction including potential resting) and “commuting” (fast and directed movement) as categories even though three-state models had lower delta AICs. The model was fitted using step lengths (assuming states could be described using a mixture of Gamma distributions), and turning angles, with wrapped Cauchy distributions. Behavioural categories were also corroborated by visual inspection after the classification.

### Foraging parameters

We calculated the straightness index for each outbound commuting flight. Tortuous commutes would indicate exploratory behaviour. Straightness was calculated by extracting the Euclidean distance between the first and last point of commutes and dividing that distance by the sum of the step lengths of the track (mean ± st dev, 1 = straight movements, 0 = tortuous movements, Table S3).

We extracted foraging events from each night and calculated the proportion of time bats spent foraging on or off Isla Colón. We tested the differences in foraging off and on Isla Colón with a binomial generalised linear model (GLM). First, we tested differences in foraging by season, using group/period and location of foraging as fixed effects. Subsequently, we tested sex differences in foraging on and off Isla Colón using group/period and sex as fixed effects. Significance threshold was p ≤ 0.05.

### Simulations and bearings

We simulated alternative tracks reflecting the movement and behavioural dynamics of the tracked bats, to estimate how these observed foraging patterns deviated from a null model given the landscape availability. We derived a three-state HMM from the initial HMM. This model included the states “foraging”, “outbound commutes” and “inbound commutes”. Commuting states were parameterized as biased correlated random walks (CRW) including angle to the colony as covariate for the mean turning angle. We restricted transitions between the two commuting states. Finally, we included the sine and cosine of time of day scaled from 0 to 1, and the square root of distance to the nearest coastline as covariates on the transition probability matrix. For the latter covariate we allowed the response to vary by including an interaction term indicating whether bats were above land, or not so that transitions from outbound commute to foraging became virtually impossible for non-land locations. We regularised data to a sampling interval of 120 s using a continuous-time CRW model (crawl) as implemented in the momentuHMM package [12], and fit separate models for seasons to account for possible differences in behaviour. Using the colony locations as starting points, we simulated alternative tracks using wet and dry season models (3 x observed tracks). We did not simulate wet season tracks for colony 3 as no observations were available. The length of the simulated tracks was chosen from a uniform distribution reflecting the interquartile range of the length of observed trajectories.

### Contrasting foraging distance and bearing between colonies and seasons

We determined foraging locations from simulated and observed tracks to compare observed and expected foraging locations. We retained only the first foraging location of each foraging bout with a duration > 0 s. We determined the proportion of foraging locations on and off Isla Colón for simulated and observed foraging locations (Figure S2). For each foraging location, we calculated the angle and distance to the colony and compared how means and variance differed between simulation and observation using a multivariate model. This was restricted to foraging locations off Isla Colón as they represented the majority of foraging.

We fitted a linear model of angles and distances to estimate the agreement between colonies and seasons for observed foraging locations (equations in Table S4), and between simulations and observed data. We included multiple observations of individuals as a random effect. We fit the model separately for each colony in the wet and dry seasons for observed and simulated data, and included weakly regularising priors.

We further computed contrasts to facilitate the evaluation of hypotheses. Contrasts - the distribution of differences between the distributions of parameters estimated by the model - were calculated to determine the difference of the population mean, effective standard deviation, and individual-level variability between wet and dry season for each colony. Contrasts were calculated as wet/dry season for angle and distance parameters. We calculated contrasts per colony and season to assess the agreement between observed and simulated foraging locations. We derived a spatial representation of the model estimates to test if similar angle and distance imply shared foraging space between colonies. We estimated the percentage of overlap between colonies during the dry season using the contours of the 2D-densities, clipped to land only. Models were implemented in STAN via CmdStan (version 2.34.1) and CmdStanR (version 0.8.1.9000).

## Results

### Mainland foraging and long-distance commutes in both dry and wet seasons

All bats with at least one completely tracked night (n = 59, colony 1: 29 (dry) - 6 (wet), colony 2: 12 (dry) - 9 (wet), colony 3: 3 (dry)) predominantly used distant foraging locations, crossing to the mainland or other islands. However, 44 bats also foraged on Isla Colon during both seasons, comprising > 30% of their total foraging (Figure S1A, GLM, p = 0.01). Females and males spent similar time foraging on and off Isla Colón (Figure S1B, p (on-island) = 0.09, p (off-island) = 0.32). Overall bats from each colony maintained long, straight commutes across seasons (Figure 1, Table S3).

Bats foraged further from the colony during the dry seasons, (Figure 2A: top left panel), with the shortest distance estimated for colony 3 and larger distances for colonies 1 and 2. Mean wet season distance was shorter in colony 1, whereas the model was inconclusive for colony 2 (see Table 1 for details).

**Table 1.** Model estimates for population means of distance and angle (estimate and 95% credibility intervals (qi)) from colonies for observed foraging locations. km = kilometres; rad = radians.

| colonies | season | mean distance [95% Qi (km)] | mean angle (rad) [95% Qi (rad)] | mean angle (degrees) [95% Qi (degrees)] |
| --- | --- | --- | --- | --- |
| colony 1 | dry | 23.47 [22.91 - 24.04] | 1.54 [1.44 - 1.65] | 268.23 [260.85 - 274.92] |
| colony 1 | wet | 18.31 [16.27 - 20.35] | 0.92 [0.5 - 1.34] | 232.7 [143.24 - 311.96] |
| colony 2 | dry | 23.23 [22.64 - 23.85] | 1.43 [1.37 - 1.49] | 261.93 [253.05 - 267.63] |
| colony 2 | wet | 21.62 [19.23 - 24.15] | 0.98 [0.4 - 1.56] | 236.15 [22.92 - 356.16] |
| colony 3 | dry | 16.05 [14.48 - 17.48] | 0.09 [-0.02 - 0.22] | 185.16 [-1.15 - 234.16] |

**Figure 2.**
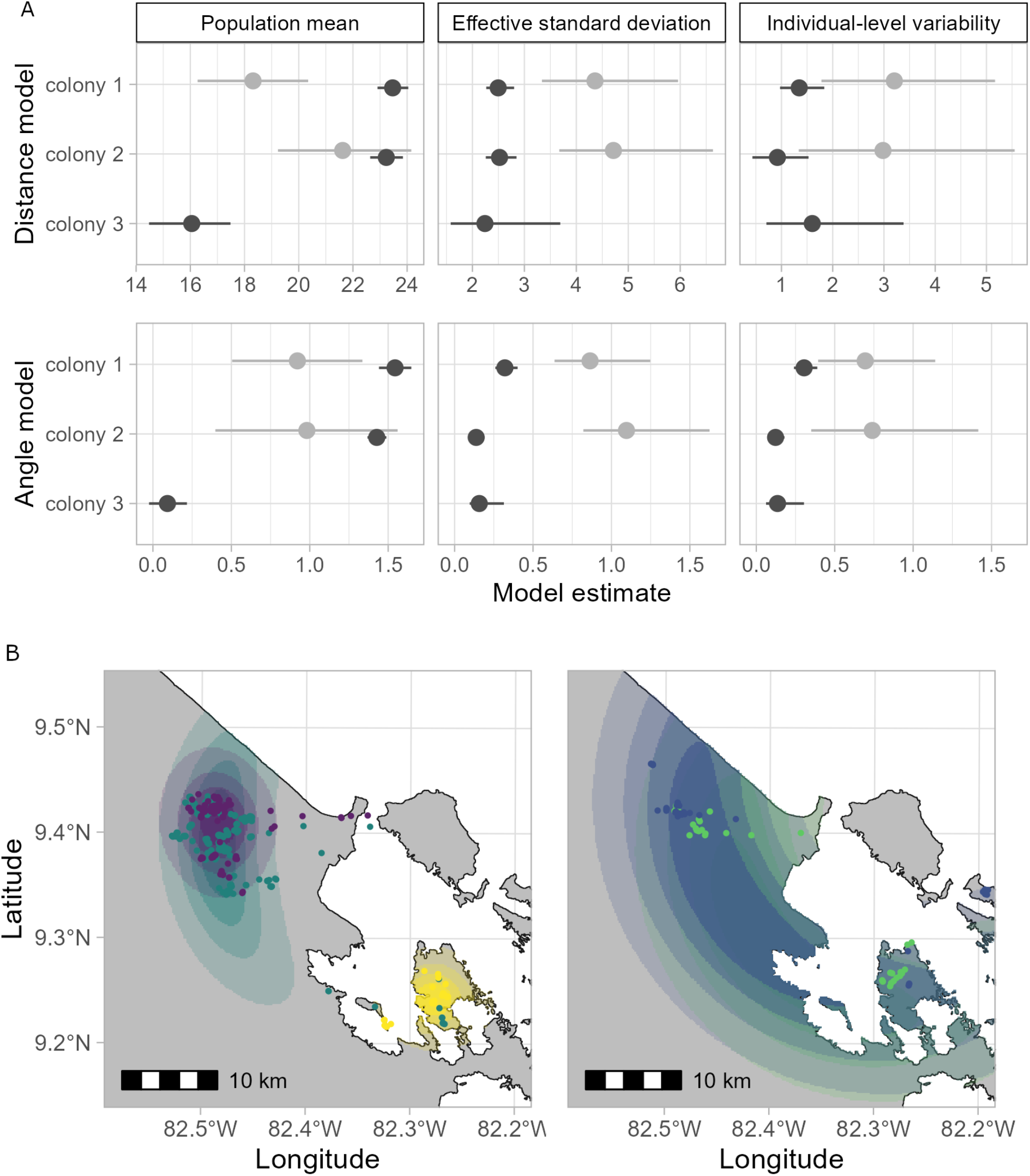
**A)** Mean and 95% credibility interval for model estimates on population means. Population mean, effective standard deviation, and individual-level variability for distance (upper row) and angle (lower row) to foraging locations. Wet season: light grey, dry season: dark grey. **B)** Spatial representation of model estimates of foraging locations beyond Isla Colón. Shown are the scaled product of the distance and angle probability density functions, clipped to the 95% contour and coastline. Colony 1: green, colony 2: blue, colony 3: yellow, intensity of colour: relative density of the PDF product.

Distances and angles from the roosting caves varied more in the wet season compared to the dry season (Figure 2). Effective standard deviation as well as the deviation of individual means from the population mean were higher during the wet season for both distance and angle (Figure 2A: centre and right panels), albeit this varied between colonies and model parameters. Differences were more pronounced for the effective standard deviation than for individual-level variability, with 95% credibility intervals showing a small level of overlap for all but the angle model in colony 2 (Figure 2A: lower right panel).

### Partial shared foraging distances, directions and space use across seasons and colonies

Individuals within the same colony, and sometimes between colonies shared foraging distances and directions. Foraging areas of colony 1 and 2 were at similar mean angles during both seasons, but colony 3, tracked only during the dry season, foraged at a much more southerly site (Figure 2B, Table 1).

The mean estimates suggested that foraging areas of colony 1 and 2 overlapped substantially but colony 3 did not: 49.58% of the area covered by colony 2 overlaps with that of colony 1, and inversely 99.17% of the area covered by colony 1 was shared with colony 2. This was indicated by contours of the probability density functions (PDFs) from distance and angle models.

### Assessing observed vs expected space use

Finally, to test our data against our predictions, we compared parameter estimates for observed and simulated foraging locations. The covariates and constraints on the transition probability matrix meant that the model was able to replicate the overall behaviour of the observed trajectories, excluding simulated foraging that fell on the ocean. While mean distance to foraging locations was similar between observations and simulations for colony 1 during the dry season (mean [95% qi]: 21.90 [20.15 - 23.59] km), simulated foraging locations were further from the colony for colonies 2 (mean [95% qi]: 29.07 [26.76 - 31.42] km) and 3 (mean [95% qi]: 37.05 [30.83 - 43.08] km) than observed locations, respectively. Simulated wet season foraging distances were longer for colony 1 (mean [95% qi]: 21.67 [18.4 - 24.72] km), but shorter than colony 2 (mean [95% qi]: 16.78 [13.2 - 20.42] km) than observed (Figure S3).

The simulations were not informed about the distribution of available resources. Thus, mean angle to the colony differed between observation and simulation, and the variance around the mean was greater for simulations (Figure 3). The models confirmed that the effective standard deviation was much greater than observed during the dry season for colony 1 (mean [95% qi]: 0.89 [0.82 - 0.97] rad), 2 (mean [95% qi]: 0.69 [0.64 - 0.74] rad), and 3 (mean [95% qi]: 1.17 [0.94 - 1.47] rad) (Figure S3). This difference was less pronounced during the wet season when there was greater spread around observed foraging locations, with effective standard deviation for simulated locations estimated as (mean [95% qi]: 0.82 [0.65 - 1.05] rad) for colony 1 and (mean [95% qi]: 1.39 [1.19 - 1.64] rad) for colony 2 (Figure S3).

**Figure 3.**
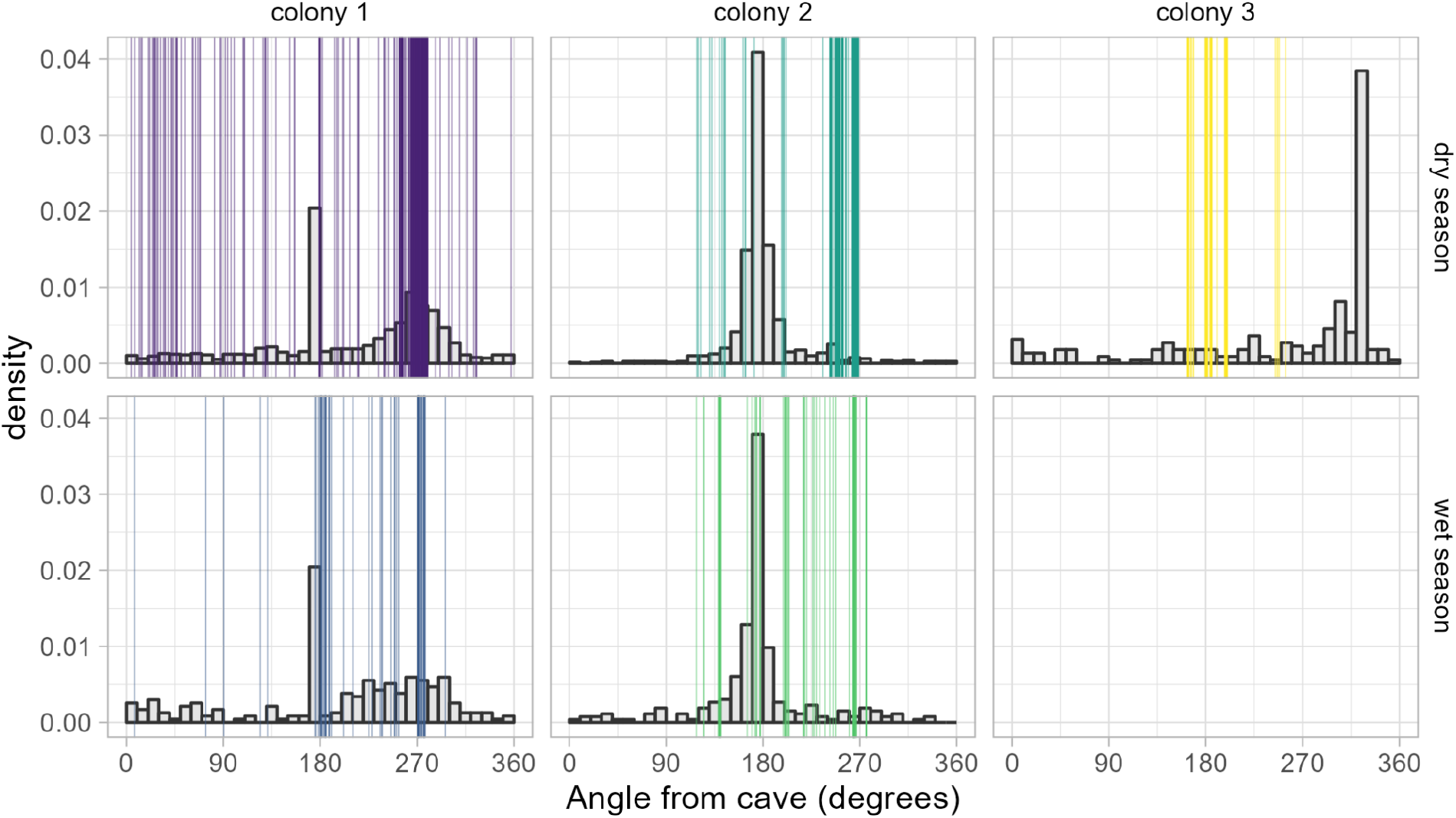
Observed and simulated angles of commute endpoints for each colony. Histogram: expected angle distribution based on simulations. Vertical lines: endpoints of outbound commutes observed in the tracking data.

## Discussion

The unique opportunity to follow foraging behaviour of the same bat species from the same island over more than six years revealed consistent colony-level behaviours across years and seasons. Bats consistently used distant foraging sites 15-25 km from the roost, much further than the <10 km previously reported [9,10]. Using distant foraging sites repeatedly can provide benefits, but the degree and profitability of this behaviour depend on the spatiotemporal predictability, quality, and depletability of a given resource [14–16]. The consistent colony-level of foraging areas across years, within seasons, and, with additional foraging areas, between seasons, suggests this behaviour could be due to familiarity [15,17]. Familiarity and the decision to exploit known foraging locations can confer long-term energetic benefits if these locations have higher productivity in temporally unpredictable environments [18]. Consistent foraging patterns help individuals to learn the location of food [19,20], move efficiently through the environment [16,21,22], or reduce conflict with neighbours [23].

Different individuals from different years but from the same colony (colony 1 and 2) used consistent foraging locations. *Phyllostomus hastatus* is highly social and capable of learning from others. It is, thus, possible that this consistency in foraging sites arises through the social transmission of information about the location of profitable resources [24,25], information use at the central place [26,27], or by following others to find unpredictable resources. In Trinidad, this species forms long-term groups of unrelated females that cooperate on multiple levels, including pup-guarding [28] and recruiting each other to feeding trees during the dry season [29]. Based on the social system described from Trinidad, we expected females to show more similar foraging patterns than males [9], but we observed no difference between the sexes. The observed long-term foraging fidelity suggests colony-level foraging preferences learned through socially transmitted information from others at the level of colony instead of female group.

We tracked colony 1 during the early and late dry season and expected increased exploration, i.e., increased path tortuosity or a win-stay, lose-shift foraging strategy [30]. With the ongoing season the switching rate to new foraging areas should match the temporal scale of resource variability (i.e., reduced balsa flower production). Instead, overall site fidelity and path straightness were maintained (Figure 1, Table S3). This does not match a change in foraging strategy linked to locally changing resources and an exploitation instead of exploration strategy. Only one individual exploited a completely different area and another exhibited exploratory behaviour (Figure S4A).

This mismatch with expectations continued when comparing seasons. Although foraging distances were shorter in the wet season when they feed on a more diverse and less ephemeral diet, bats still mostly foraged off Isla Colón. We had also expected less shared foraging space in the wet season. Although some individuals switched foraging areas they were still in a shared direction, perhaps in a mix of win-stay, lose-shift foraging in learned preferred foraging areas (Figure S4B). This is confirmed by the bats continuing to show little exploration, but directed and straight commuting flights. Our results indicate that during part of the year, *P. hastatus* may switch between a set of socially learned foraging areas, rarely individually exploring the landscape for food.

The long foraging distances in 2016 were thought to be due to unusually late balsa flowering [11]. Thus, the continued long foraging distances over the years, when balsa as well as more ubiquitously distributed wet season resources should have been available on the island were particularly surprising (Figure 1, 3). The use of shared foraging areas is likely a choice rather than a fixed behaviour. Bats spent up to 60-100 minutes a night commuting, time and effort they could have spent feeding or exploring closer to the roost, avoiding the risk of crossing open water. Why they continue to invest time and energy to travel to these distant foraging areas remains unresolved, but is likely based on some learned traditions as they are clearly able to visit and use closer resources (Figure S2).

Individuals from the same colonies consistently used shared foraging distances and directions. This differed somewhat between colonies. We had expected this due to competition for limited balsa flowers during the dry season [11]. It is interesting that colony 2 used similar foraging areas to colony 1, even though colony 3 is geographically closer. One possibility is dispersal of knowledgeable individuals between colonies transferred information that spread through the colony. Additionally, shared foraging areas could indicate particularly high balsa tree availability, enough to sustain at least two colonies of 500 bats each [31]. At peak flower production, a balsa tree can feed 3-7 bats over one night [11]. Thus 72 -166 trees would be needed to satisfy the energetic demands of one of these colonies. Ground truthing indicated high balsa availability in these areas, and future studies should incorporate measures of flower availability. Overall, the continued use of a similar area even in the wet season when feeding on insects and fruit, reinforces the idea that the use of foraging areas is acquired through memory and possibly conformity, rather than density-dependent or between colony competition, as observed in other frugivorous bats, such as *Rousettus aegyptiacus* [20].

Additional, non-exclusive aspects may play a role in the use of shared foraging areas. Bats tracked during 2016 did not share flowering trees, but rested together between foraging bouts, potentially to exchange information or increase vigilance against predators [11]. Resolution of tracking data after 2016 was lower to increase the duration of data collection. This made it impossible to test for social resting but we confirmed that bats returned to the same foraging patches within the shared general foraging area night after night.

Our results indicate strong colony foraging preferences that are independent of seasonality and group composition. However, these results represent only a partial picture of the wide range of behavioural strategies that *P. hastatus* might have. Two main limitations remain unresolved: our inability to track bats for long-term periods and our lack of detailed knowledge of *P. hastatus* diet and resource availability for a species that moves tens of kilometres. Our research usually assumes that animal behaviour is always completely adaptive, but our results suggest that animals can choose foraging behaviours that do not follow the predictions of ideal foraging and optimising returns for reasons we have yet to understand.

## Ethics

This study was conducted under the permit of Ministerio del Ambiente, Panama (SE/A-96-15, SE/A-96-18, SE/A-38-2020), and the Animal Care and Use Committee at the Smithsonian Tropical Research Institute (2014-0701-2017, 2017-0815-2020-A2, 2020-0212-2023), and adhered to the ASAB/ABS Guidelines for the Use of Animals in Research.

## Data and Code availability

All data and code are available at www.movebank.org and https://github.com/mccalderonc/P_hastatus_Consistentlong-distance_foragingflights

## Declaration of AI use

We have not used AI-assisted technologies in creating this article.

## Authors’ contributions

MCCC, MTO, MCC, DKND: conceive the manuscript. MCCC, MTO, TDB, MCC, DKND: collected biologging data. MCCC, MvT: did the analyses. MCCC, MvT, MTO, MCC, DKND: wrote the manuscript.

## Acknowledgements

We thank the Smithsonian Tropical Research Institute (STRI) especially Plinio Gondola and Urania Gonzales. We are also grateful to the local people who allowed us to work on their properties. Special thanks to the STRI bat lab for their support. Cynthia Peña, Edward Hurme, Frederic Touzalin, Graciela Aguilar, Jann Sarapak, James Lee, Lucia Torrez, Luisa Gomez, Michael Spiedel, Nina Hwang, Ricardo Cossio, Richard Emlet, Tadhg Lonergan and especially Leslye Barria helped during data collection.

