## Supplementary material for "Consistent long-distance foraging flights across years and seasons at colony level in a Neotropical bat": Figure S1

**Figure S1. Proportion of time bats spent foraging on and off the island extracted from the behavioural classification from the Hidden Markov Models (HMM).** A) Bats spent most of the time foraging off Isla Colón during both seasons (less than 30% on Isla Colón). B) Foraging off and on Isla Colón was similar for females (right panel) and males (left panel). dark grey: foraging off island, light grey: foraging on Isla Colón. Error bars: standard error from the mean values.

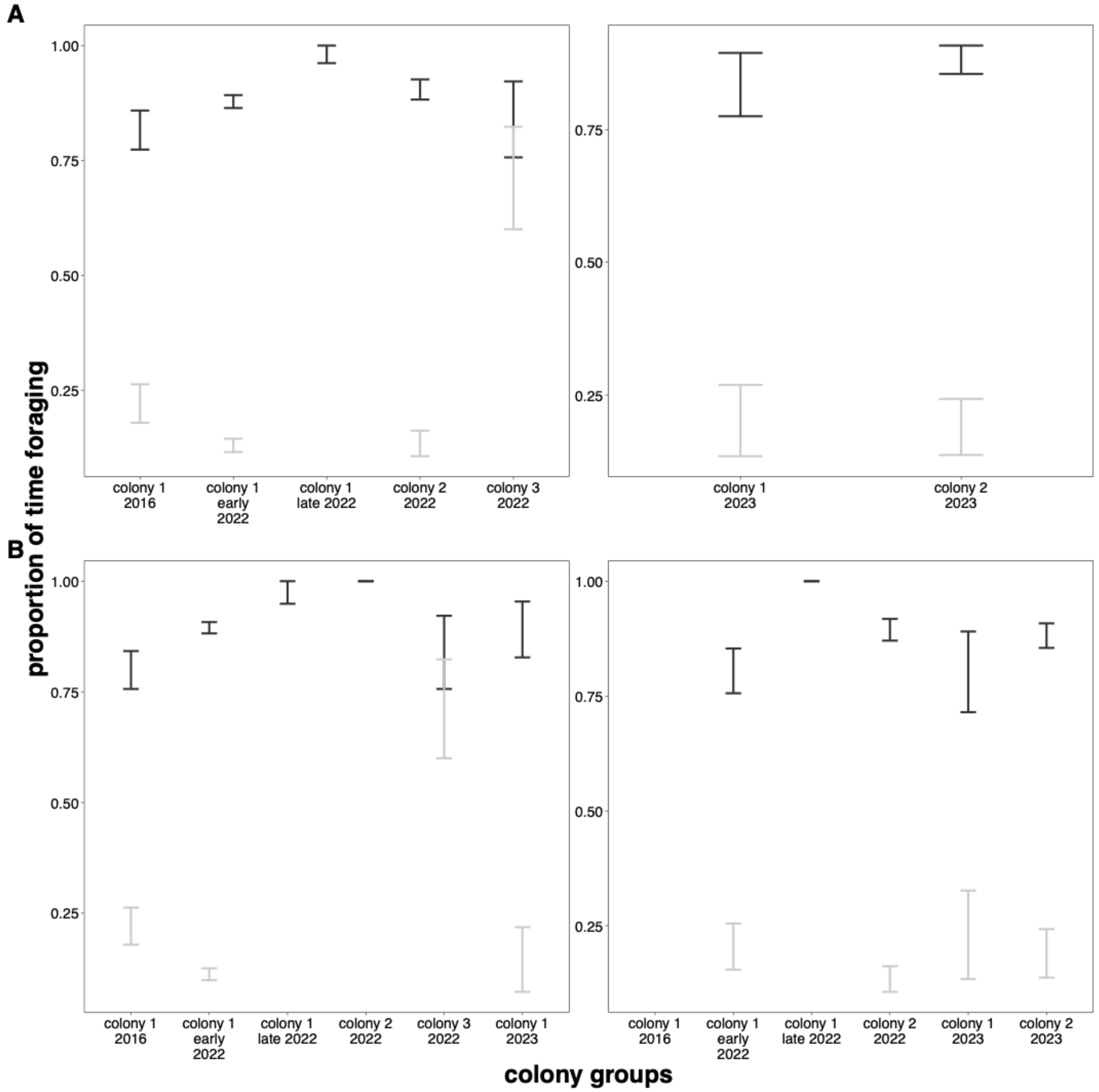
