## Supplementary material for "Consistent long-distance foraging flights across years and seasons at colony level in a Neotropical bat": Figure S2

**Figure S2.** Proportion of foraging locations on and off Isla Colón for observed and simulated foraging locations.

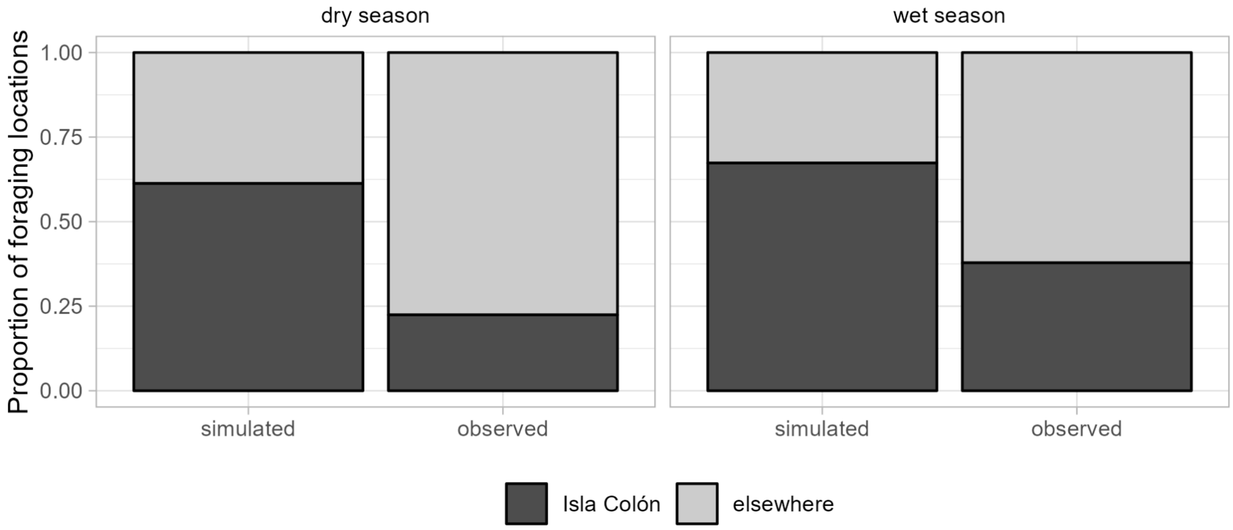
