## Supplementary material for "Consistent long-distance foraging flights across years and seasons at colony level in a Neotropical bat": Figure S3

**Figure S3. Model estimates for the simulated data.** Mean and 95% credibility interval for model estimates from the simulated data on population mean, effective standard deviation, and individual-level variability for distance (upper row) and angle (lower row) to foraging locations. Wet season: light grey, dry season: dark grey.

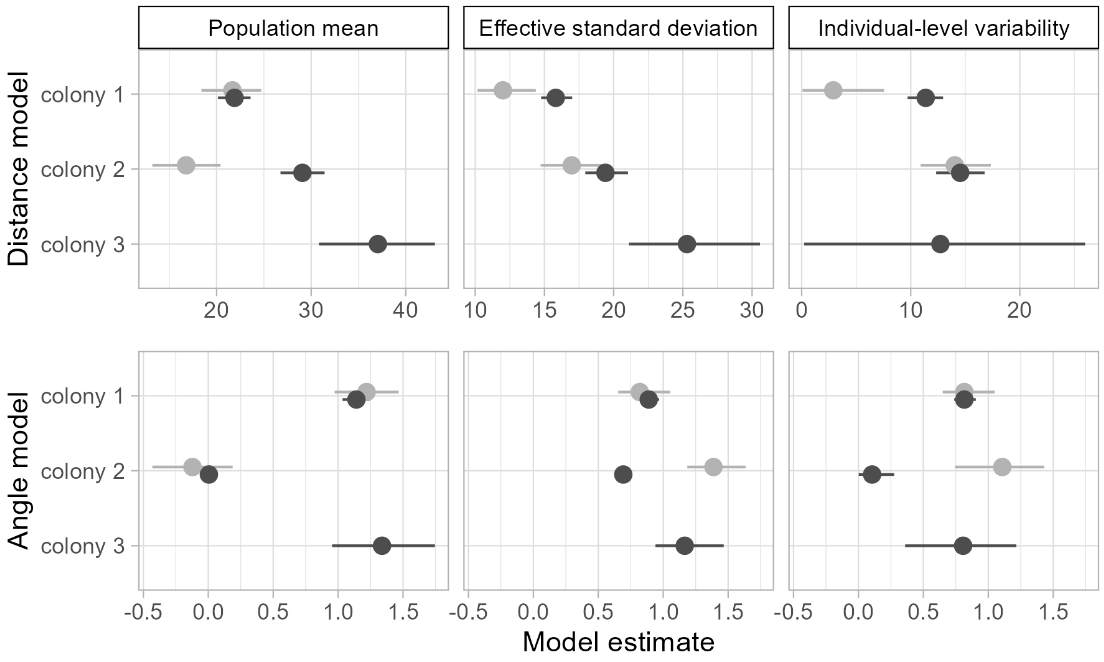
