## Supplementary material for "Consistent long-distance foraging flights across years and seasons at colony level in a Neotropical bat": Figure S4

**Figure S4**. Foraging area switching and exploratory behaviour. A) During the dry season, one individual in colony 1 exhibited exploratory behaviour (top) and another one used a different foraging area (bottom). B) During the wet season three individuals from colony 2 switched foraging areas.

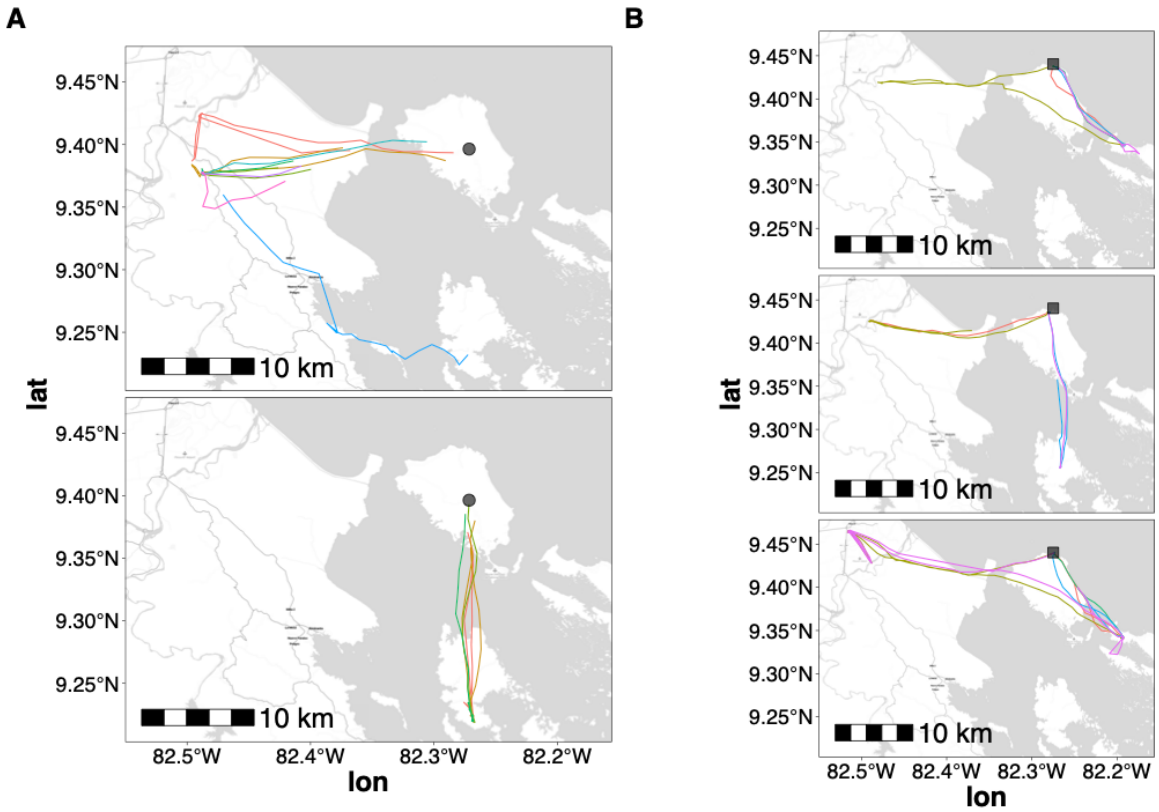
