## Supplementary material for "Consistent long-distance foraging flights across years and seasons at colony level in a Neotropical bat": Table S1

**Table S1.** Information on tagged bats and tag programming schedules. Mass is given in grams, repro (i.e., reproductive status) in females is nulli = nulliparous/has never reproduced; preg = pregnant; lac = lactating; plac = post-lactating; in males: nr = non-reproductive or scr = scrotal/testes enlarged; colonies are 1-3, 2016 data from [11]; dry = bat was tagged during the dry season, wet = bat was tagged during the wet season, early = bat tagged in February 2022 and late = bat tagged in March 2022. Details on programming schedules: In 2016 tags went into a low energy sleep state for five minutes and then restarted to search for satellites for 90 s, when GPS reception was not available. Tag performance varied depending on how deep in the cave bats were roosting and the density of foliage where bats perched while foraging. In March 2022 tags went to sleep for 10 min and searched for satellites for 15 s, when no GPS reception was available. In August 2023 tags also collected continuous ACC data at 25Hz with 4G, 10 bits.

| bat Id | bat mass | repro | sex | tag brand | GPS fix frequency (secs) | tag mass | colony | year | season |
|---|---|---|---|---|---|---|---|---|---|
| 2016030703 | 137 | nr | m | Gypsy-5 | 1 | 7 | 1 | 2016 | dry |
| 2016030705 | 118 | plac | f | Gypsy-5 | 1 | 7 | 1 | 2016 | dry |
| 71A0D95 | 122 | nulli | f | Gypsy-5 | 1 | 7.5 | 1 | 2016 | dry |
| 71A111A | 109 | nulli | f | Gypsy-5 | 1 | 7.25 | 1 | 2016 | dry |
| 74D8954 | 128 | plac | f | Gypsy-5 | 1 | 6.5 | 1 | 2016 | dry |
| 74D8C25 | 131 | nulli | f | Gypsy-5 | 1 | 7 | 1 | 2016 | dry |
| 74D932E | 116 | nulli | f | Gypsy-5 | 1 | 7 | 1 | 2016 | dry |
| 74D972D | 122 | plac | f | Gypsy-5 | 1 | 7 | 1 | 2016 | dry |
| 74DA035 | 126 | nulli | f | Gypsy-5 | 0.5 | 6.5 | 1 | 2016 | dry |
| 74DA92F | 109 | plac | f | Gypsy-5 | 1 | 6.5 | 1 | 2016 | dry |
| 74DAF9C | 121 | plac | f | Vesper | 1 | 6.5 | 1 | 2016 | dry |
| 74DC454 | 116 | plac | f | Vesper | 1 | 6.5 | 1 | 2016 | dry |
| 74DCA83 | 123 | nulli | f | Gypsy-5 | 1 | 6.5 | 1 | 2016 | dry |
| 74DCBCC | 114 | plac | f | Gypsy-5 | 1 | 7 | 1 | 2016 | dry |
| 74DDA80 | 123 | nulli | f | Gypsy-5 | 1 | 7.25 | 1 | 2016 | dry |
| 74DDFB1 | 124 | plac | f | Vesper | 1 | 7 | 1 | 2016 | dry |
| 74DE4E7 | 124 | plac | f | Gypsy-5 | 0.5 | 7 | 1 | 2016 | dry |
| 74DE9A7 | 122 | plac | f | Gypsy-5 | 1 | 7 | 1 | 2016 | dry |
| 74F7D4C | 137 | nr | m | Gypsy-5 | 1 | 7 | 1 | 2016 | dry |
| 74F8E19 | 113 | plac | f | Gypsy-5 | 0.5 | 6 | 1 | 2016 | dry |

| | | | | | | | | | |
|---|---|---|---|---|---|---|---|---|---|
| 74F9F83 | 116 | plac | f | Gypsy-5 | 0.5 | 6 | 1 | 2016 | dry |
| 74FE24E | 113 | plac | f | Gypsy-5 | 0.5 | 6 | 1 | 2016 | dry |
| 0C506E35_C | 134 | nr | m | Vesper | 30 | 8.45 | 1 | 2021 | wet |
| 0D501435_C | 119 | lac | f | Vesper | 30 | 8.52 | 1 | 2021 | wet |
| 31507235_C | 95 | lac | f | Vesper | 30 | 6.11 | 1 | 2021 | wet |
| 32501535_C | 121 | nr | m | Vesper | 30 | 8.67 | 1 | 2021 | wet |
| 38500337_C | 91 | lac | f | Vesper | 30 | 6.04 | 1 | 2021 | wet |
| 38506D37_C | 99 | lac | f | Vesper | 30 | 8.52 | 1 | 2021 | wet |
| 53507135_C | 131 | lac | f | Vesper | 30 | 8.71 | 1 | 2021 | wet |
| 5C500E4E_C | 119 | nr | m | Vesper | 30 | 8.59 | 1 | 2021 | wet |
| 0C506E35_G | 110 | plac | f | Vesper | 120 | 6 | 1 | 2022 | early dry |
| 23506B4E_G | 118 | plac | f | Vesper | 120 | 6 | 1 | 2022 | early dry |
| 2C501A35_G | 110 | plac | f | Vesper | 120 | 6 | 1 | 2022 | early dry |
| 2C507735_G | 111 | nulli | f | Vesper | 120 | 6 | 1 | 2022 | early dry |
| 2D507235_G | 114 | nulli | f | Vesper | 120 | 6 | 1 | 2022 | early dry |
| 2E500235_G | 111 | plac | f | Vesper | 120 | 6 | 1 | 2022 | early dry |
| 31507235_G | 120 | plac | f | Vesper | 120 | 6 | 1 | 2022 | early dry |
| 38506737_G | 140 | nr | m | Vesper | 120 | 5.5 | 1 | 2022 | early dry |
| 39506037_G | 115 | nulli | f | Vesper | 120 | 6 | 1 | 2022 | early dry |
| 53506935_G | 120 | plac | f | Vesper | 120 | 6 | 1 | 2022 | early dry |
| 22507B4E_D | 120 | plac | f | Vesper | 120 | 5.88 | 1 | 2022 | early dry |
| 2C500235_D | 132 | nr | m | Vesper | 120 | 6.36 | 1 | 2022 | early dry |
| 37506237_D | 102 | lac | f | Vesper | 120 | 6.29 | 1 | 2022 | early dry |
| 37507537_D | 112 | lac | f | Vesper | 120 | 6.19 | 1 | 2022 | early dry |
| 38500437_D | 131 | nr | m | Vesper | 120 | 6.12 | 1 | 2022 | early dry |
| 38506D37_D | 138 | nr | m | Vesper | 120 | 5.88 | 1 | 2022 | early dry |
| 39500E37_D | 138 | plac | f | Vesper | 120 | 5.89 | 1 | 2022 | early dry |
| PH_TS_004 | 91 | nulli | f | Axy-track | 180 | 7 | 2 | 2022 | early dry |
| PH_TS_011 | 130 | nr | m | Axy-track | 180 | 7 | 2 | 2022 | early dry |
| PH_TS_014 | 122 | nr | m | Axy-track | 180 | 7 | 2 | 2022 | early dry |
| PH_TS_016 | 130 | nr | m | Axy-track | 180 | NA | 2 | 2022 | early dry |
| PH_TS_018 | 119 | preg | f | Axy-track | 180 | 6 | 1 | 2022 | late dry |
| PH_TS_024 | 143 | nr | m | Axy-track | 180 | 7 | 2 | 2022 | early dry |
| PH_TS_029 | 119 | preg | f | Axy-track | 180 | 6 | 3 | 2022 | early dry |
| PH_TS_030 | 126 | scr | m | Axy-track | 180 | 7 | 1 | 2022 | late dry |
| PH_TS_039 | 117 | nr | m | Axy-track | 180 | 7 | 1 | 2022 | late dry |
| PH_TS_046 | 121 | preg | f | Axy-track | 180 | 7 | 1 | 2022 | late dry |
| PH_TS_049 | 129 | NA | f | Axy-track | 180 | 7 | 3 | 2022 | early dry |
| PH_TS_052 | 126 | nr | m | Axy-track | 180 | 6 | 3 | 2022 | early dry |
| PH_TS_056 | 128 | pre | f | Axy-track | 180 | 6 | 3 | 2022 | early dry |

| | | | | | | | | | |
|---|---|---|---|---|---|---|---|---|---|
| PH_TS_062 | 128 | plac | f | Axy-track | 180 | 6 | 3 | 2022 | early dry |
| PH_TS_072 | 114 | preg | f | Axy-track | 180 | 6 | 3 | 2022 | early dry |
| PH_TS_074 | 131 | nr | m | Axy-track | 180 | NA | 2 | 2022 | early dry |
| PH_TS_079 | 135 | nr | m | Axy-track | 180 | 7 | 2 | 2022 | early dry |
| PH_TS_080 | 136 | nr | m | Axy-track | 180 | 7 | 2 | 2022 | early dry |
| PH_TS_081 | 110 | nulli | f | Axy-track | 180 | 7 | 2 | 2022 | early dry |
| PH_TS_083 | 128 | nr | m | Axy-track | 180 | 7 | 2 | 2022 | early dry |
| PH_TS_085 | 128 | nr | m | Axy-track | 180 | 7 | 2 | 2022 | early dry |
| PH_TS_096 | 101 | nulli | f | Axy-track | 180 | 7 | 1 | 2022 | late dry |
| PH_TS_098 | 119 | nr | m | Axy-track | 180 | 7 | 2 | 2022 | early dry |
| PH_TS_100 | 117 | nr | m | Axy-track | 180 | 7 | 2 | 2022 | early dry |
| PH_TS_103 | 116 | preg | f | Axy-track | 180 | 7 | 2 | 2022 | early dry |
| PH_TS_112 | 133 | nr | m | Axy-track | 180 | 7 | 2 | 2022 | early dry |
| PH_TS_113 | 131 | nr | m | Axy-track | 180 | 7 | 2 | 2022 | early dry |
| PH_TS_114 | 131 | nr | m | Axy-track | 180 | 7 | 2 | 2022 | early dry |
| PH_TS_117 | 140 | preg | f | Axy-track | 180 | 7 | 1 | 2022 | late dry |
| PH_TS_120 | 123 | nr | m | Axy-track | 180 | 7 | 2 | 2022 | early dry |
| PH_TS_121 | 114 | nr | m | Axy-track | 180 | 7 | 2 | 2022 | early dry |
| PHYL1 | 126 | nr | m | Axy-track | 60 | 4.8 | 1 | 2023 | wet |
| PHYL11 | 136 | nr | m | Axy-track | 60 | 4.8 | 2 | 2023 | wet |
| PHYL16 | 146 | nr | m | Axy-track | 60 | 4.8 | 2 | 2023 | wet |
| PHYL21 | 180 | nr | m | Axy-track | 60 | 4.8 | 2 | 2023 | wet |
| PHYL22 | 142 | nr | m | Axy-track | 60 | 4.8 | 2 | 2023 | wet |
| PHYL24 | 126 | nr | m | Axy-track | 60 | 4.8 | 2 | 2023 | wet |
| PHYL25 | 172 | nr | m | Axy-track | 60 | 4.8 | 2 | 2023 | wet |
| PHYL27 | 127 | nr | m | Axy-track | 60 | 4.8 | 2 | 2023 | wet |
| PHYL28 | 137 | nr | m | Axy-track | 60 | 4.8 | 2 | 2023 | wet |
| PHYL29 | 123 | nulli | f | Axy-track | 60 | 4.2 | 1 | 2023 | wet |
| PHYL3 | 112 | nulli | f | Axy-track | 60 | 4.8 | 1 | 2023 | wet |
| PHYL30 | 113 | nulli | f | Axy-track | 60 | 4.2 | 1 | 2023 | wet |
| PHYL31 | 111 | nulli | f | Axy-track | 60 | 4.2 | 1 | 2023 | wet |
| PHYL32 | 118 | nulli | f | Axy-track | 60 | 4.2 | 1 | 2023 | wet |
| PHYL33 | 126 | nulli | f | Axy-track | 60 | 4.2 | 1 | 2023 | wet |
| PHYL34 | 110 | nulli | f | Axy-track | 60 | 4.2 | 1 | 2023 | wet |
| PHYL35 | 146 | nr | m | Axy-track | 60 | 4.2 | 1 | 2023 | wet |
| PHYL36 | 128 | nulli | f | Axy-track | 60 | 4.2 | 1 | 2023 | wet |
| PHYL37 | 110 | nulli | f | Axy-track | 60 | 4.2 | 1 | 2023 | wet |
| PHYL38 | 117 | nulli | f | Axy-track | 60 | 4.2 | 1 | 2023 | wet |
| PHYL39 | 150 | nr | m | Axy-track | 60 | 4.2 | 1 | 2023 | wet |
| PHYL4 | 135 | nr | m | Axy-track | 60 | 4.8 | 1 | 2023 | wet |

| | | | | | | | | | |
|---|---|---|---|---|---|---|---|---|---|
| PHYL40 | 112 | nulli | f | Axy-track | 60 | 4.2 | 1 | 2023 | wet |
| PHYL7 | 173 | nr | m | Axy-track | 60 | 4.8 | 2 | 2023 | wet |
| PHYL9 | 170 | nr | m | Axy-track | 60 | 4.8 | 2 | 2023 | wet |
