## Supplementary material for "Consistent long-distance foraging flights across years and seasons at colony level in a Neotropical bat": Table S2

**Table S2.** Summary table of the number of individuals used within a colony/period for the analyses.

| colony | year | season | N tags recovered | N individuals | N females | N males |
|---|---|---|---|---|---|---|
| colony 1 | 2016 | dry | 22 | 16 | 15 | 1 |
| colony 1 | 2022 | early dry | 17 | 16 | 12 | 4 |
| colony 1 | 2022 | late dry | 6 | 5 | 4 | 1 |
| colony 2 | 2022 | dry | 19 | 18 | 3 | 15 |
| colony 3 | 2022 | dry | 6 | 6 | 5 | 1 |
| colony 1 | 2021 | wet | 8 | 2 | 2 | 0 |
| colony 1 | 2023 | wet | 15 | 12 | 8 | 4 |
| colony 2 | 2023 | wet | 10 | 10 | 0 | 10 |
