## Supplementary material for "Consistent long-distance foraging flights across years and seasons at colony level in a Neotropical bat": Table S3

**Table S3.** Mean and standard deviation of straightness index for all colonies and periods during both dry and wet seasons. The index values range from zero to one. 0 = tortuous movements, 1 = straight movements.

| **Colonies** | **Straightness index** | |
|---|---|---|
| | **dry** | **wet** |
| Colony 1 | 0.93±0.1 (2016)<br>0.88±0.16 (early 2022)<br>0.84±0.16 (late 2022) | 0.93±0.01 (2021)<br>0.90±0.09 (2023) |
| Colony 2 | 0.76±0.23 (2022) | 0.90±0.07 (2023) |
| Colony 3 | 0.81±0.23 (2022) | |
