## Supplementary material for "Consistent long-distance foraging flights across years and seasons at colony level in a Neotropical bat": Table S4

**Table S4.** Equations used in the linear model for distance and angles for the observed and simulated data. The effective standard deviations capture the overall variability, for each colony and season.

| **Equation for distance** | **Parameter explanation** |
|---|---|
| $distance_i \sim Normal(\mu_{distance[i]}, \sigma)$<br>$\mu_i \sim Normal(M_{distance}, \tau)$<br>$M \sim Normal(0,5)$<br>$\sigma \sim Exponential(1)$<br>$\tau \sim Exponential(0.5)$, | distance = distance of the *i*th observation<br>$\mu_i$ = mean distance of the *i*th observation<br>M = mean distance of all observations<br>$\tau$ = deviation of the individual mean distances from M (the individual-level variability)<br>$\sigma$ = standard deviation of the distances (the observation-level variability). |
| **Equation for angle** | |
| $angle_i \sim Normal(\mu_{angle[i]}, \kappa)$<br>$\mu_{angle[i]} \sim Normal(M_{angle}, \zeta)$<br>$M_{angle} \sim Normal(0,1)$<br>$\kappa \sim Exponential(1)$<br>$\zeta \sim Exponential(0.5)$, | $\kappa$ = individual-level variability<br>$\zeta$ = standard deviation of the angle to foraging locations<br>$\sqrt{\tau^2 + \sigma^2}$ = effective standard deviation from mean population distances<br>$\sqrt{\kappa^2 + \zeta^2}$ = effective standard deviation from mean population angles |
